# Multi-omics characterization of astrocyte subtypes reveals spatially coordinated astrocyte downregulation in depression

**DOI:** 10.64898/2026.07.02.735636

**Authors:** Haruka Mitsuhashi, Harish R. Rao, Steffanie Amadei, Anjali Chawla, Maria Antonietta Davoli, Naguib Mechawar, Gustavo Turecki, Corina Nagy

## Abstract

Major depressive disorder (MDD) is a complex psychiatric disorder affecting millions of individuals worldwide. Astrocytes, which have been implicated in MDD by several studies, are the most abundant non-neuronal cells in the brain and play critical roles in synaptic regulation, blood-brain barrier maintenance, and immune modulation. While astrocytic molecular and morphological abnormalities are well-established features of MDD, these alterations have not been resolved within their spatial context. Here, we combine spatial transcriptomics with matched snRNA-seq and snATAC-seq datasets to spatially map molecularly distinct astrocyte subtypes and define their regional contributions to MDD pathology. This spatial context further enables the characterization of astrocyte interactions with neighboring cell populations, providing a more holistic assessment of how dysfunctional astrocytes influence local brain microenvironments and circuit function in MDD. We identified spatially localized astrocytic dysfunction in deep cortical layers of the MDD dlPFC, converging across transcriptomic, chromatin, and spatial modalities and centering on the PSAP-GPR37L1 signaling axis. Together, these findings identify astrocyte dysfunction as a key feature of MDD and demonstrate the value of spatially resolved molecular profiling for uncovering how altered astrocyte–neuron communication within deep cortical layers may contribute to disease pathology.

## Introduction

Major depressive disorder (MDD) is a complex psychiatric disorder affecting approximately 300 million people worldwide ^1^. Recent advances in cell-type-specific profiling have provided important insights into its molecular pathology. Using single-nucleus RNA sequencing (snRNA-seq), we previously identified transcriptional dysregulation in deep-layer excitatory neurons and several glial populations, including oligodendrocyte precursor cells, microglia, and astrocytes ^2, 3^. Complementary single-nucleus ATAC sequencing (snATAC-seq) analyses revealed alterations in chromatin accessibility, particularly in deep-layer excitatory neurons and microglia, accompanied by differential transcription factor motif accessibility ^4^.

Astrocytes play essential roles in maintaining brain function through synaptic regulation, blood-brain barrier integrity, and immune modulation ^5^. Disruption of these processes has been implicated in MDD pathology ^6^. Emerging evidence further suggests that astrocytes are key mediators of the stress response, as they are highly responsive to glucocorticoid signalling ^7, 8^. Transcriptomic studies have consistently showed downregulation of astrocyte-specific genes in both postmortem human brain and animal models of depression ^9, 10^. Likewise, numerous studies show morphological changes to distinct astrocytic subtypes in MDD ^11-13^. Together, these findings suggest that chronic stress may contribute to astrocyte dysfunction, supporting a central role for astrocytes in MDD pathology.

One of the critical roles of astrocytes is to support neurons and other glial cells through spatially organized interactions ^14, 15^. However, previous approaches lack spatial context, limiting our ability to associate transcriptional changes to cell-cell communication. Spatial transcriptomics addresses this limitation by enabling gene expression measurement while preserving the spatial location of transcripts. Recent studies applying spatial transcriptomics to the human prefrontal cortex of healthy individuals have successfully mapped the laminar organization of the dlPFC, providing insight into spatially resolved gene expression patterns ^16, 17^.

Here, we characterized astrocyte subtypes in the human dlPFC and investigate their dysregulation in MDD through an integrative multi-omics approach. We applied spatial transcriptomics to postmortem dlPFC tissue from 16 individuals. By integrating these data with matched snRNA-seq and snATAC-seq datasets from overlapping subjects, we mapped astrocyte subtypes across cortical layers and characterized their molecular profiles ^2-4^. We identified layer-specific transcriptomic alterations, with astrocyte dysfunction in deep cortical layers. Furthermore, spatially organized cell-cell communication programs were characterized by astrocytic-specific functions, with the alterations to the main molecular players being supported by complementary data.

## Results

### Characterization of Astrocytic subtypes

To characterize astrocytes in the dlPFC, we analyzed previously published snRNA and snATAC datasets, which have 94% samples overlap. The snRNA dataset comprised 71 subjects, including 34 psychiatrically healthy controls and 37 individuals diagnosed with MDD (Figure 1A; Supplementary Figure 1) ^2, 3^. The dataset contained a total of 160,811 nuclei, of which 13,117 were annotated as astrocytes and classified into two subtypes. The snATAC data included additional subjects (n=84 subjects), comprised 40 psychiatrically healthy controls and 44 individuals diagnosed with MDD, and generated approximately 200,000 nuclei, of which 13,610 were identified as astrocytes ^4^. Four astrocyte subclusters were identified in the original clustering, comprising 6,437 nuclei in Ast1, 2,318 in Ast2, 3,426 in Ast3, and 1,429 in Ast4 ^4^. Cis-regulatory elements (CREs) in each subtype were characterized in the previously published data ^4^.

**Figure 1:**
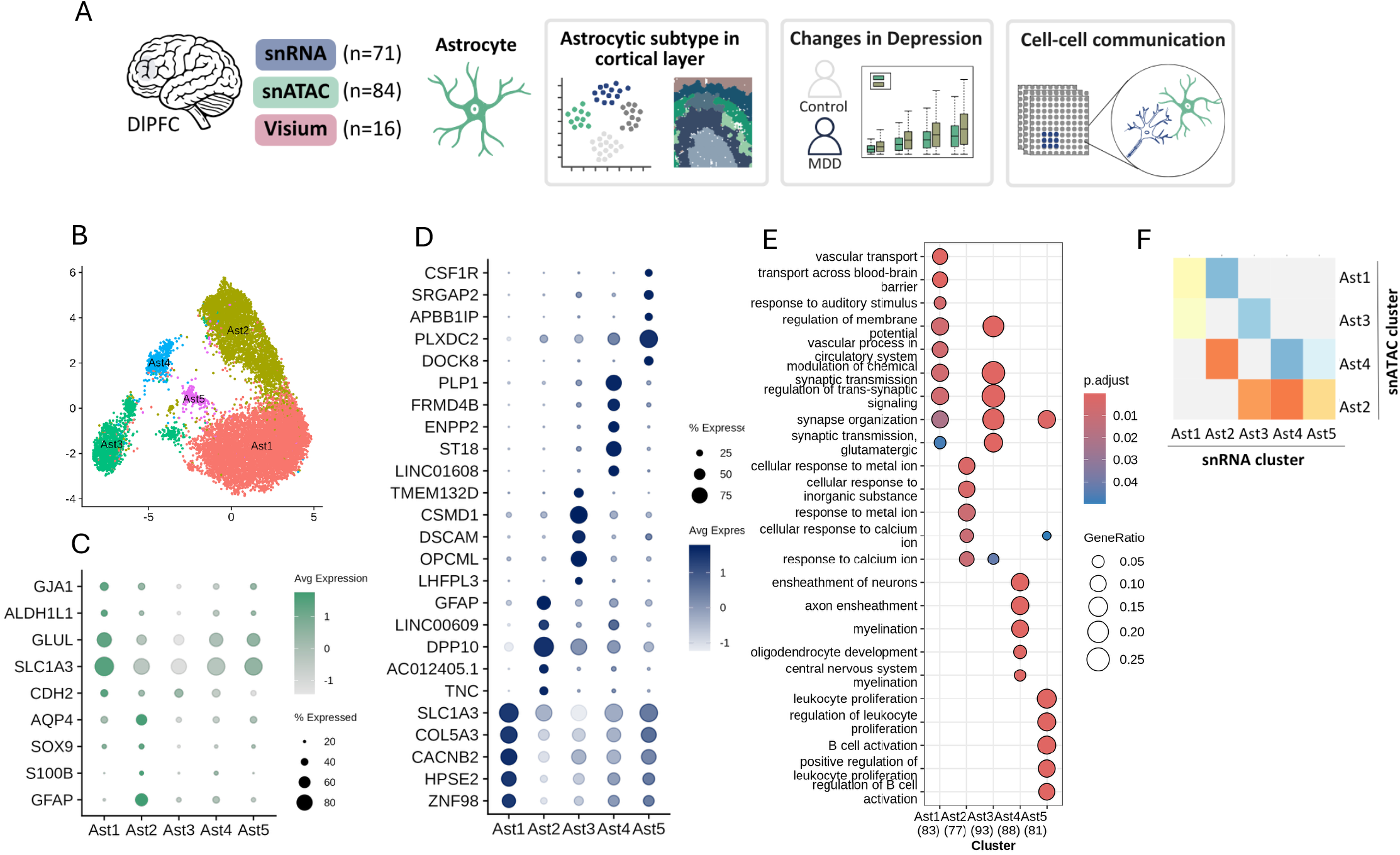
Characterization of astrocyte subtypes using the snRNA-seq data. **A**. Schematic of the study design integrating snRNA-seq, snATAC-seq, and spatial transcriptomics data. Multiomic datasets including snRNA-seq (n = 71), snATAC-seq (n = 84), and Visium spatial transcriptomics (n = 16) from human dlPFC were integrated to identify cortical layer-associated astrocyte subtypes, characterize depression-associated molecular alterations, and investigate changes in cell-cell communication networks in MDD. **B**. UMAP plot of astrocyte subclusters identified following re-clustering of the astrocyte population in snRNA-seq data, colored by subtype. **C**. Gene expression of selected astrocyte marker genes across subtypes. **D**. Gene expression of the top five marker genes per astrocyte subtype. **E**. Gene ontology analysis performed using the top 100 marker genes per astrocyte subtype, showing distinct functional enrichment across subtypes. **F**. MetaNeighbor heatmap showing correspondence between astrocyte subclusters identified in snRNA-seq (rows) and astrocyte subtypes identified in snATAC-seq (columns), with color representing AUROC score.

As shown by astrocytic subtypes in the snATAC data, cortical astrocytes are known to be heterogeneous populations ^3^. This led us to further resolve astrocyte nuclei in the snRNA data. We performed sub-clustering using Seurat and identified five populations of astrocyte (Figure 1B). Among these, Ast1 and Ast2 represented the largest populations, comprising 6,966 and 3,804 nuclei, respectively. Ast1 showed higher expression of genes involved in glutamate metabolism (GLUL, SLC1A3) and gap junction protein (GJA1), whereas Ast2 expressed higher levels of reactive astrocyte markers (GFAP, S100B), supporting a functional distinction of these astrocytes (Figure 1C). Gene ontology analysis of Ast1 revealed enrichment for vascular, blood–brain barrier, and synaptic-related processes, suggesting interactions with both the neurovascular unit and neuronal populations (Figure 1E). The smaller clusters exhibited mixed signatures. Ast3 expressed neuronal-associated genes (DSCAM, OPCML) and was enriched for synaptic signaling pathways (Figure 1D and 1E). Ast4 displayed a mixed glial signature, characterized by gene enrichment of myelination-related terms. Ast5 was characterized by immune-related genes (CSF1R, APBB1IP, DOCK8), consistent with enrichment of leukocyte- and B-cell-associated pathways.

To assess molecular similarities between astrocyte subclusters across the snRNA and snATAC datasets, we used MetaNeighbor, which quantifies the reproducibility of cell type identities across independent datasets ^18^. Ast2 in the snRNA-seq data corresponded most closely to Ast4 in the snATAC-seq data (AUROC = 0.87; Figure 1F). In contrast, Ast1 in the snRNA-seq data showed moderate similarity to two snATAC-seq clusters, Ast1 and Ast3 (AUROC = 0.68 and 0.61, respectively). The three smaller snRNA clusters (Ast3, Ast4, and Ast5) showed the highest similarity to Ast2 in the snATAC-seq data, suggesting that these populations share a common transcriptional and chromatin accessibility state across modalities.

### Concordant transcriptional and chromatin accessibility changes

Next, we investigated the association of astrocyte subtype-specific transcriptomics changes with MDD. A total of 43 genes were identified using the same threshold as in the original analysis ^3^ (adj. p-value < 0.1 and |logFC| > log2(1.1)), 31 gene in Ast1 and 12 genes in Ast2 (Figure 2A; Supplementary Table 4). Most of these genes were downregulated in MDD, 90 % and 83 % genes respectively. No significant DEGs were identified in Ast3, Ast4, and Ast5, and therefore, subsequent analyses focused exclusively on Ast1 and Ast2.

**Figure 2:**
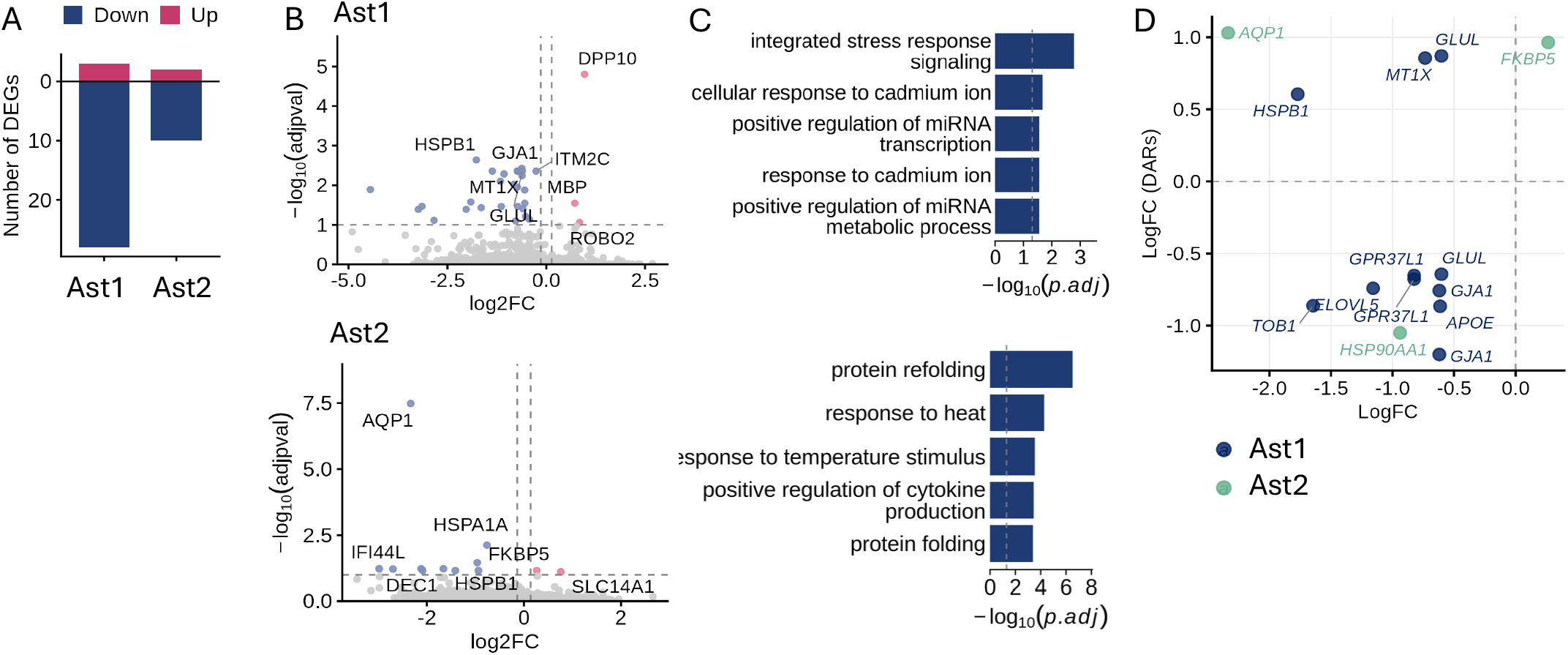
Differentially expressed genes in astrocyte subtypes in the snRNA-seq data. **A**. Number of DEGs identified in each astrocyte subtype in MDD compared to controls, colored by direction of fold change (downregulated = blue, upregulated = red), using a threshold of adjusted p-value < 0.1 and |logFC| > log2(1.1). **B**. Volcano plot of DEG results in each astrocyte subtypes, with top DEGs labeled, colored by direction of fold change. **C**. Gene ontology enrichment analysis of downregulated DEGs, showing the top five enriched biological processes per subtype. No significant enrichment was identified in upregulated genes. **D**. Scatter plot showing overlap between DEGs and DAR-linked genes (adjusted p-value < 0.2 and |logFC| > log2(1.1)), with log fold change of gene expression on the x-axis and log fold change of chromatin accessibility on the y-axis. Genes appearing multiple times reflect links to more than one differentially accessible region. Points are colored by astrocyte subtype.

In Ast1, genes known to have astrocyte specific functions were found to be differentially expressed, for instance, genes involved in gap junction (GJA1), synaptic function/plasticity (GLUL, AGT), lipid metabolism (APOE, CLU, ANGPTL4, ELOVL5, CSTS), water transport (AQP4) were identified (Figure 2B). Gene ontology analysis showed that DEGs in Ast1 were enriched for stress response signaling pathways whereas Ast2 were enriched for response to steroid hormone (Figure 2C). Interestingly, the downregulation in both subtypes is driven by males, as sex-stratified differential expression analyses identified higher number of DEGs in males compared with females (Supplementary Figure 2D).

We then assessed concordance between gene expression and chromatin accessibility changes. Across the matched astrocyte subtypes identified above, we compared DEGs with the previously identified DAR-linked genes (*r*>0.45) using the original threshold ^4^ (adj. p-value < 0.2 and |logFC| > log2(1.1)). In Ast1, 8 genes were identified as both DEGs and linked to DARs, the majority of which were downregulated in MDD (73%), implicating decreased chromatin accessibility driving reduced gene expression (Figure 2D). These included key astrocyte genes, GLUL (DEG: adj. p-value = 0.0044, DAR: adj. p-value = 0.096, 0.169), GJA1 (DEG: adj. p-value = 0.0037, DAR: adj. p-value = 0.0224, 0.113), APOE (DEG: adj. p-value = 0.0057, DAR: adj. p-value = 0.0623), GPR37L1 (DEG: adj. p-value = 0.0095, DAR: adj. p-value =0.091, 0.169). In Ast2, 3 concordant DEG-DAR pairs were identified. FKBP5 showed increased chromatin accessibility alongside increased gene expression in MDD (DEG: adj. p-value = 0.0697, DAR: adj. p-value =0.197), while AQP1 showed increased chromatin accessibility but decreased gene expression (DEG: adj. p-value = 3.28E-08, DAR: adj. p-value =0.172).

We then asked whether there was a difference in transcription factor binding site accessibility in either of these astrocytic subpopulations that might regulate these effects. Using ChromVar, we previously reported that the NR3C1 motif for the glucocorticoid receptor was more accessible in MDD, specifically in Ast1 (adj. p-value = 7.28 × 10^−5^; Supplementary Figure 3) ^4^. This finding is consistent with the enrichment of stress-response signaling pathways in Ast1 DEGs, suggesting that altered glucocorticoid receptor activity may contribute to the transcriptional changes.

### Cortical layer structures using the spatial transcriptomics

To further characterize astrocyte subtype heterogeneity and dysfunction within their native tissue context, we generated spatial transcriptomic maps of the dorsolateral prefrontal cortex (dlPFC) from 16 subjects using the 10x Genomics Visium platform, including 8 psychiatrically healthy controls and 8 individuals diagnosed with MDD. Unsupervised clustering was performed using Banksy to identify cortical layer structures (Figure 3B; Supplementary Figure 10B) ^19^. The identified spatial domains which were annotated and validated against a histological annotated reference dataset ^16, 20^ (Figure 3C and 3D; Refer to Methods). Seven spatial domains corresponding to cortical layers were identified as WM, WM/L6, L6, L5, L4, L3, and L2.

**Figure 3:**
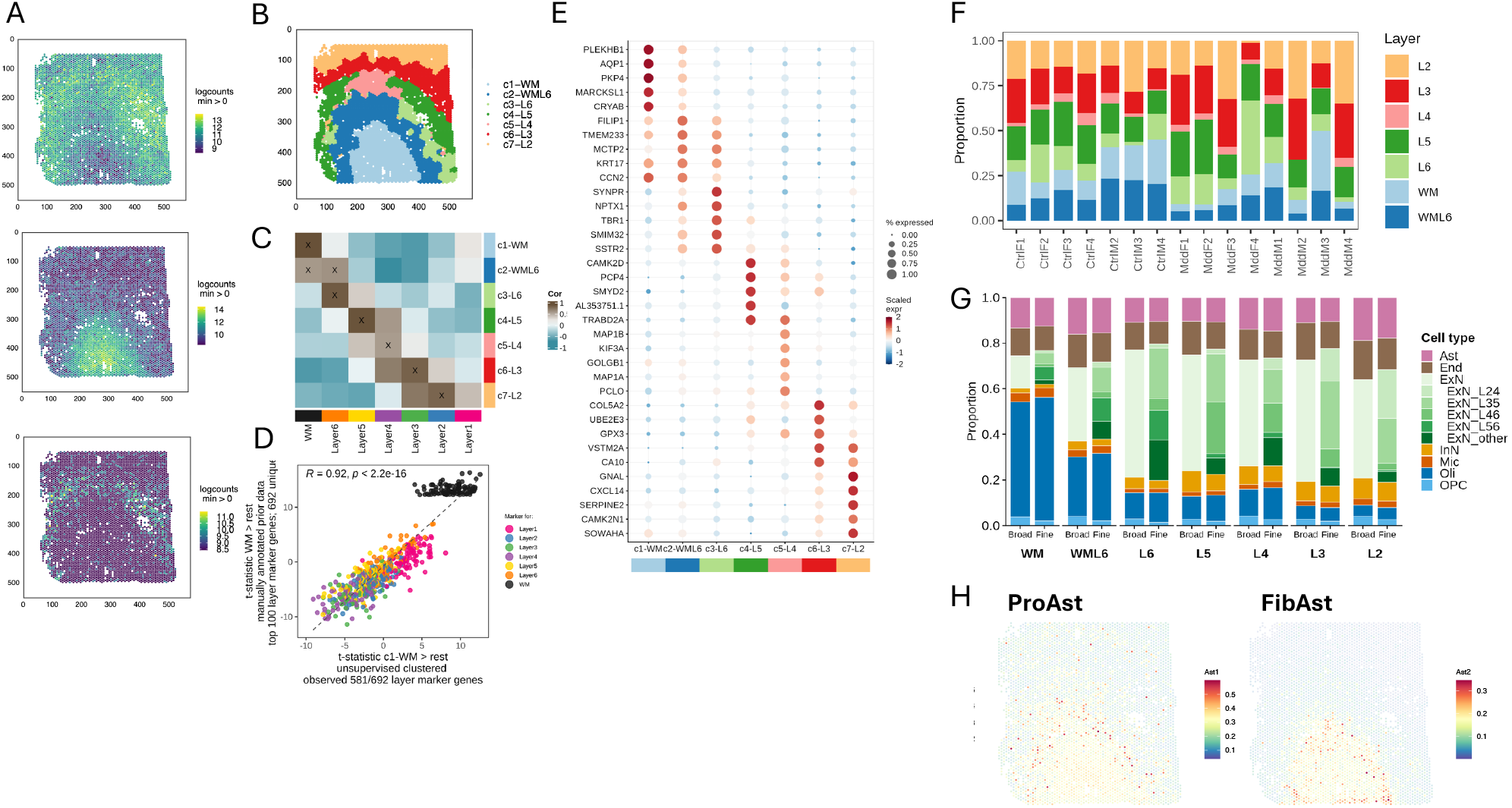
Characterization of cortical layers in the dlPFC using the Visium data. **A**. Feature plots showing expression of neuronal marker NeuN, astrocyte marker GFAP, and deep-layer marker PCP4 in a representative sample, reflecting the laminar organization of the dlPFC. **B**. Spatial domains identified by unsupervised clustering using Banksy at resolution 0.6, colored by identified spatial domains. **C**. Heatmap showing correlation of enrichment scores between our identified spatial domains (columns) and manually annotated dlPFC Visium data from the reference data (rows), colored by correlation coefficient. X marks indicating confidently matched domains (correlation > 0.25, merge ratio = 0.1). **D**. Scatter plot of enrichment t-statistics comparing white matter against all other layers in the manually annotated reference data (x-axis) versus our identified spatial domains (y-axis), demonstrating high concordance between datasets. A correlation of (R = 0.92, p < 2.2 × 10^−16^) showing high agreement with the reference dataset. **E**. Gene expression of top 5 genes identified as a gene marker in each spatial domain. **F**. Proportion of spots assigned to each cortical layer (L1–L6) and white matter (WM) for each subject. F and M denote female and male subjects, respectively. Colors indicate layer identity. **G**. Mean cell type proportion per cortical layer estimated by Cell2location spot deconvolution, colored by cell type. Deconvolution was performed at two levels: broad cell types and fine-level excitatory neuron subtypes with layer-specific identity. **H**. Spatial distribution of snRNA-seq-defined astrocyte subtypes mapped onto Visium spots using label transfer. Ast1 was enriched in deep cortical layers and displayed a protoplasmic astrocyte phenotype (ProAst; left), whereas Ast2 was enriched in white matter and displayed a fibrous astrocyte phenotype (FibAst; right).

Due to the resolution of the 10x Visium platform, each spot captures transcripts from multiple cells. To estimate the cell type composition, we performed spot deconvolution using Cell2location ^21^. Spot deconvolution was performed at two levels, a broad cell type level and a fine-level annotation of excitatory neuron subtypes with layer identity. Consistent with the literature, oligodendrocytes were strongly enriched in the c1–WM domain (46.1%), while astrocytes were distributed across cortical layers (10.8–17.8%) (Figure 3G). Excitatory neuron subtypes showed high enrichment across cortical layers. Using the fine-level annotation, excitatory neuronal subtypes showed the highest enrichment in their corresponding cortical layers, with ExN_L46 enriched in L5 (21.8%), ExN_L23 in L3 (28%), and ExN_L24 in L2 (20.3%).

Next, we sought to characterize the spatial distribution of the astrocyte subtypes that showed molecular changes in MDD. snRNA subtypes labels were transferred to spatial transcriptomics data using Seurat label transfer ^22^. Ast2 identified in the snRNA-seq data was enriched in white matter, consistent with a fibrous astrocyte phenotype that is typically localized to this region ^23^ (Figure 3H). In contrast, Ast1 was enriched in cortical layers, consistent with a form of protoplasmic astrocytes. For clarity, we hereafter refer to Ast1 as protoplasmic astrocytes and Ast2 as fibrous astrocytes.

### Concordant downregulation of astrocyte activities across modalities

We next examined layer-specific transcriptional changes associated with MDD in the spatial transcriptomic data. Using the threshold (adj. p-value < 0.1 and |FC| > 1.5), a total of 28 DEGs were identified across cortical layers (Figure 4A; Supplementary Table 6). The highest number of DEGs was observed in Layer 6, of these, 13 genes were downregulated and 1 gene was upregulated in MDD relative to controls.

**Figure 4:**
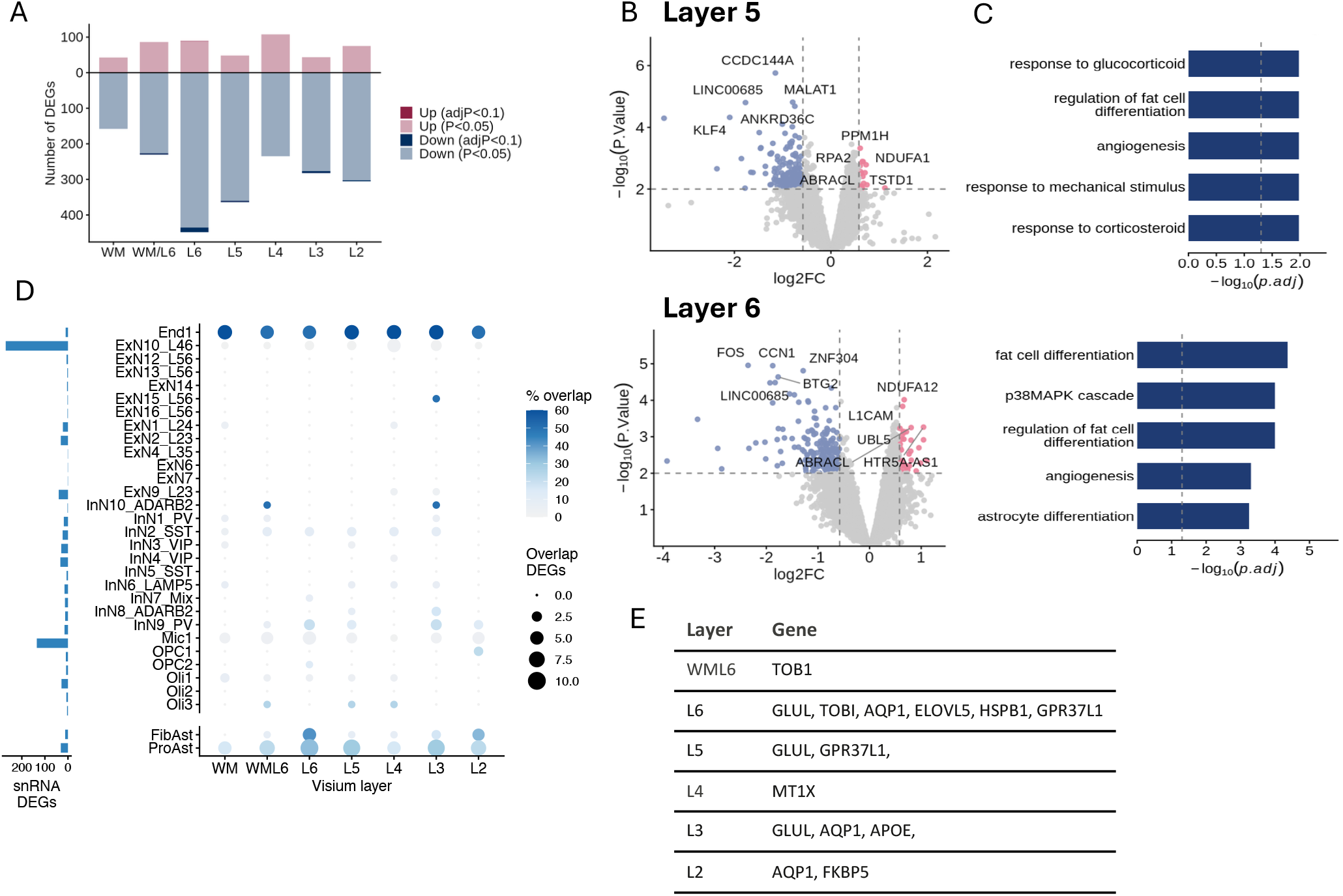
Differentially expressed genes in each cortical layer in the Visium data. **A**. Number of DEGs identified in each spatial domain in MDD compared to controls, colored by direction of fold change (downregulated = blue, upregulated = red). Dark colors indicate DEGs identified using a stringent threshold (adjusted p-value < 0.1), while light colors indicate DEGs identified using a lenient threshold (nominal p-value < 0.05). **B**. Volcano plot of DEG results in Layer5 (top) and Layer6 (down). Top DEGs labeled and colored by direction of fold change. **C**. Gene ontology enrichment analysis of downregulated DEGs, showing the top five enriched biological processes in Layer5 (top) and Layer6 (down). **D**. Dot plot showing the overlap between snRNA-seq DEGs and Visium layer-specific DEGs. DEGs identified in protoplasmic astrocytes, fibrous astrocytes, and other cell types from previous studies are shown separately. Dot size indicates the number of overlapping genes between each cell type and cortical layer. Color represents the percentage of snRNA-seq DEGs overlapping with layer specific DEGs. The bar plot on the left shows the total number of DEGs identified in each cell type in snRNA-seq. **E**. Table showing genes with concordant dysregulation across three modalities, DEGs identified in snRNA-seq (adjusted p-value < 0.2 and |logFC| > log2(1.1)), DEGs identified in Visium (p-value < 0.05 and |logFC| > 0.58)), and genes linked to DARs in snATAC-seq (adjusted p-value < 0.2 and |logFC| > log2(1.1)). Rows are organized by cortical layer, and all the genes indicated are downregulated in spatial transcriptomics data.

To explore convergence of molecular signals across modalities, a more lenient threshold was used (p-value < 0.05 and |FC| > 1.5). This identified 2,515 DEGs across cortical layers (Figure 4A; Supplementary Table 6), with Layer 6 consistently showing the highest number of DEGs (523 genes; Figure 4B). Of these, 435 genes (83%) were downregulated in MDD and were enriched for pathways related to angiogenesis and astrocyte differentiation (Figure 4C). Similarly, 408 DEGs were identified in Layer 5, the majority of which were downregulated (88%; Figure 4B). Downregulated genes in Layer 5 were enriched for pathways associated with glucocorticoid response and angiogenesis (Figure 4C). Overall, the downregulation of astrocyte-related genes in deep cortical layers were consistent with the downregulation of astrocytes observed in the snRNA data.

We next assessed the overlap between layer-specific DEGs and cell type-specific DEGs in the snRNA data (Figure 4D). Protoplasmic astrocytes showed the strongest concordance, with 18 of 31 DEGs (58%) downregulated in both modalities. These genes were distributed across cortical layers, with the greatest overlap observed in Layer 6 (10 genes). Fibrous astrocytes also showed substantial overlap, with 7 of 23 DEGs (30%) shared between modalities. Among non-astrocyte cell types, Mic1 microglia and ExN10_L46 neurons exhibited relatively large numbers of overlapping genes (13 and 12 genes, respectively). However, these represented a smaller proportion of their respective DEG sets, and the direction of differential expression was less consistent between modalities.

We further explored whether these differences were accompanied by chromatin accessibility changes. 9 genes showed dysregulation across all three modalities and cortical layers (Figure 4E). The largest overlap was observed in layer 6, where GLUL (p-value = 0.0029), TOB1 (p-value = 0.0082), ELOVL5 (p-value = 0.0161), HSPB1 (p-value = 0.0297), and GPR37L1 (p-value = 0.0307) from protoplasmic astrocytes showed concordant downregulation across all modalities. GLUL (p-value = 0.0188) and GPR37L1 (p-value = 0.0290) were also downregulated in layer 5. Together, these findings demonstrate concordant dysregulation of astrocyte-specific genes at both the transcriptomic and chromatin accessibility levels in MDD, with the strongest evidence observed in the deep cortical layers.

### Spatially coordinated cell-cell communication driven by astrocyte

Next, to better understand the consequences of repressed astrocytic function, we examined cell–cell communication across cortical layers using LIANA+. Specifically, we identified spatially co-localized ligand–receptor, which reflects local cell-cell communication, and applied non-negative matrix factorization (NMF) to identify coordinated signaling programs across slides (Figure 5A) ^24^.

**Figure 5:**
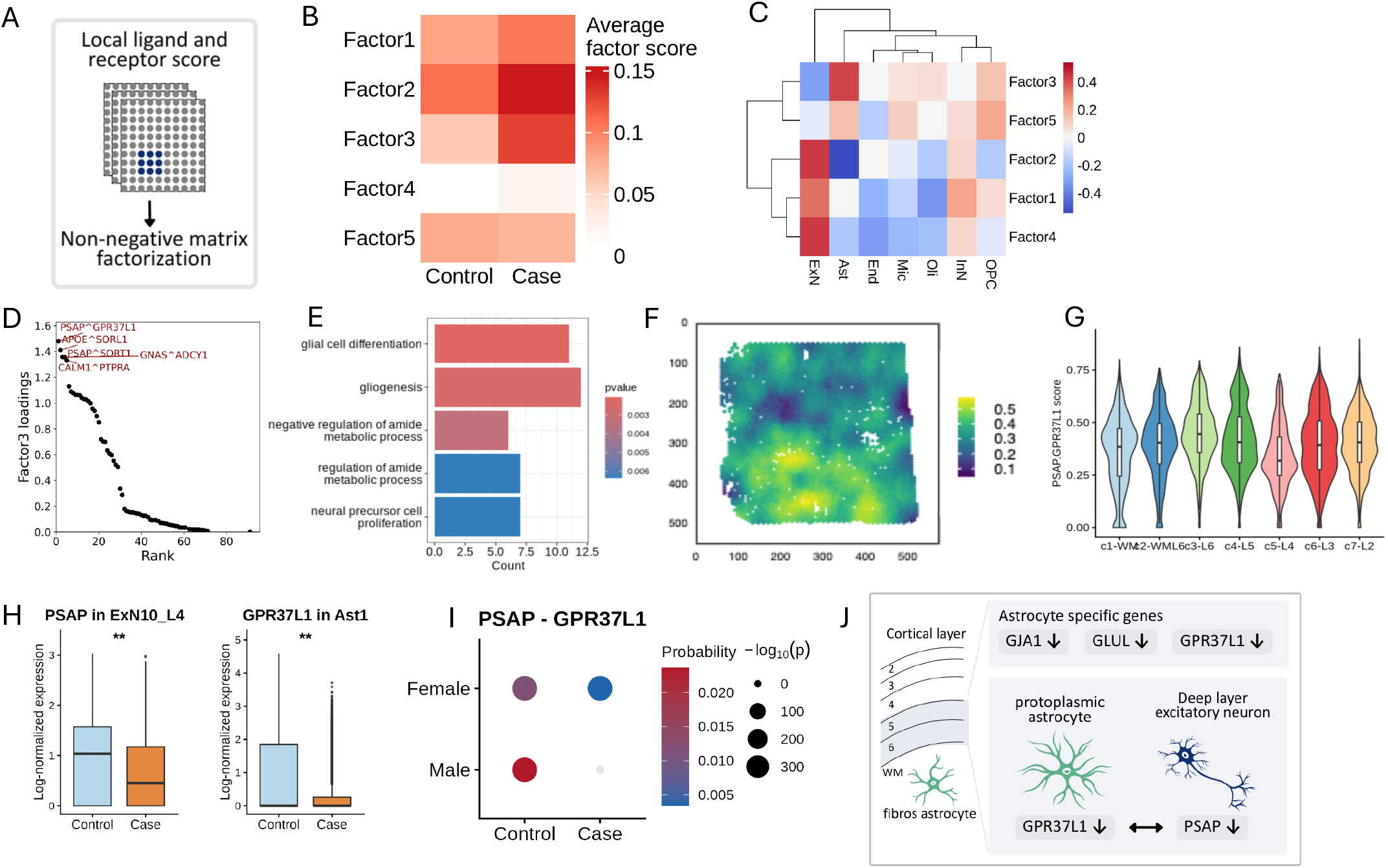
Astrocyte driven ligand and receptor communication in the Visium and the snRNA-seq data. **A**. Schematic illustrating the ligand-receptor co-expression program analysis using LIANA+. Spatially weighted ligand-receptor co-expression scores are computed per spot, followed by non-negative matrix factorization (NMF) to identify spatially organized signaling programs. **B**. Heatmap showing the average NMF factor scores across control and MDD subjects. Statistical comparison between groups were performed using Mann-Whitney U tests on subject-level average factor scores. **C**. Heatmap showing correlation between cell type proportions estimated by Cell2location and ligand-receptor factor scores across Visium spots. Cell type proportion using broad cell type annotation was used. Rows represent cell types and columns represent factors, colored by correlation coefficient. **D**. Ranked contribution of ligand-receptor pairs to Factor 3 loading scores, showing the top 5 contributing pairs. **E**. Gene ontology enrichment analysis of the top contributing ligands and receptors (standard deviation threshold = 1) in Factor 3. **F**. Spatial feature plots showing local co-expression scores of PSAP and GPR37L1 across spots. **G**. Distribution of local co-expression of PSAP and GPR37L1 across cortical layer. **H**. Bar plots showing expression levels of PSAP in ExN10_L46 excitatory neurons, GPR37L1 in Ast1 astrocytes between controls and MDD cases in the snRNA-seq data. (** adjusted p-value < 0.01). **I**. PSAP–GPR37L1 signaling between ExN10_L4/6 excitatory neurons and Ast1 astrocytes inferred from snRNA-seq data. Communication probability was estimated separately in male and female datasets, with ExN10_L4/6 excitatory neurons designated as sender cells and Ast1 astrocytes as receiver cells. Dot color represents the inferred communication probability. Dot size reflects the statistical significance of the interaction. The absence of a dot indicates that no significant communication was detected. **J**. Schematic summary of dysregulated astrocyte signaling. Protoplasmic astrocytes located in deep cortical layers exhibit downregulation of astrocyte-specific genes and reduced PSAP–GPR37L1 signaling, suggesting impaired astrocyte-mediated with excitatory neurons in deep layers.

Using this approach, we identified five ligand–receptor programs (factors), with each factor characterized by a distinct sets of ligands and receptors (Supplementary Figure 11A). Among these, Factor 4 was significantly enriched in MDD relative to controls (Mann-Whitney U test, p-value = 0.0321; Figure 5B). Gene ontology analysis of the top contributing ligands and receptors revealed enrichment for neuronal processes (Supplementary Figure 11C). Consistent with these functional annotations, Factor 4 scores were positively associated with excitatory neuron proportion in each spot (Figure 5C). However, only one gene, L1CAM, among the top contributing ligands and receptors was differentially modified in the complementary analysis.

This limited molecular overlap prompted us to examine additional communication programs associated with MDD, revealing Factor 3 as a prominent astrocyte-related signaling network (Figure 5E). Factor 3 was among the most highly enriched factors positively associated with MDD, although it only trended toward statistical significance (Mann-Whitney U test, p-value = 0.0922; Figure 5B). Key ligands and receptors contributing to this factor included GPR37L1, PTPRZ1, APOE, and SORT1, all of which are predominantly expressed in astrocytes (Figure 5D). Furthermore, 7 of the top contributing ligands and receptors (A2M, APOE, CHL1, CNTN2, FGFR3, GPR37L1, and L1CAM) were also identified as layer-specific DEGs, supporting dysregulated astrocyte-driven signaling in MDD.

### Disrupted PSAP-GPR37L1 signaling in MDD

The top-ranked ligand-receptor pair in the astrocytic associated Factor 3, Prosaposin (PSAP) and G-protein-coupled receptor 37-like 1 (GPR37L1), showed convergent evidence of dysregulation across all three modalities. GPR37L1 showed reduced expression (adj. p-value = 0.0095, logFC = -0.8245; Figure 5H) and decreased chromatin accessibility (adj. p-value =0.091, logFC = -0.677; adj. p-value = 0.169, logFC = - 0.652) in protoplasmic astrocytes as well as decreased gene expression in deep cortical layers (Layer5: p-value = 0.0290, logFC = -0.8037; Layer6: p-value = 0.0307, logFC = -0.6313). In line with this, the distribution of local PSAP–GPR37L1 signaling scores differed significantly across cortical layers (Kruskal–Wallis test, p-value = 3.09 × 10^−230^), with the highest median scores observed in Layer 6 (Figure 5F and 5G; Supplementary Figure 11E and 11F).

Consistent with known PSAP-GPR37L1 interaction by neurons and astrocytes, PSAP was also downregulated in deep layer excitatory neurons, ExN10_L46, in our snRNA-data (Figure 5H) ^3^. To assess whether this interaction is altered in MDD, we performed cell-cell communication analysis in the snRNA data using CellChat ^25^. Because the original snRNA datasets were generated separately for males and females, analyses were conducted independently within each sex to account for batch effects. In both sexes, the communication probability of PSAP-GPR37L1 signaling from ExN10_L46 neurons to Ast1 astrocytes was decreased in MDD, further supporting cell-type-specific dysregulation of PSAP-GPR37L1 signaling in MDD (Figure 5I).

## Discussion

Using converging evidence from multiple modalities, we demonstrate coordinated astrocyte dysfunction in the dlPFC of individuals with MDD. We observed downregulation of key astrocyte genes, including GLUL and GPR37L1, particularly within deep cortical layers. Spatially organized ligand-receptor co-expression programs further support disrupted astrocyte driven communication as a key feature of MDD pathology (Figure 5J).

Astrocytes have traditionally been classified into subtypes based on their morphology and anatomical localization ^26^. Recent advances in single-cell and spatial transcriptomic technologies have highlighted substantial astrocyte heterogeneity, identifying distinct populations across cortical layers and brain regions ^27-29^. By integrating snRNA-seq and spatial transcriptomic data, we identified astrocyte populations corresponding to the classical protoplasmic and fibrous astrocyte states. Consistent with previous reports, the protoplasmic astrocyte exhibited lower GFAP expression than the fibrous astrocyte ^30, 31^. Functional enrichment analysis further suggested that protoplasmic astrocytes are specialized for synaptic transmission, glutamate buffering, and vascular maintenance, indicating a synaptic-supportive protoplasmic astrocyte. Importantly, this population showed stronger transcriptional alterations in MDD. Consistent with previous studies reporting astrocyte dysfunction in MDD, genes highly specific to astrocytes were downregulated ^10, 12, 32-34^. Gene ontology analysis of the downregulated genes in protoplasmic astrocytes revealed enrichment for stress-response signaling, consistent with the known vulnerability of astrocytes to stress and glucocorticoid signaling ^7, 35^. Among the DEGs, GLUL and GJA1 was previously found to be strongly decreased in a large cohort of postmortem human dlPFC samples from individuals with MDD who died by suicide (n = 121), strongly supporting our current findings ^10^.

Here we found that the astrocyte Gap junction, Connexin-43, GJA1, showed reduced expression and decreased chromatin accessibility in MDD. Connexin-43 (Cx43) mediates intercellular communication among astrocytes and maintains astrocytic networks ^36^. Previous studies from our group ^10, 34, 37, 38^, and others ^33^, reported decreased expression of GJA1 in multiple brain regions, with these changes being partly mediated by epigenetic mechanisms. Consistent with our findings, animal models of depression have shown reduced GJA1 expression ^6^, with restoration of Cx43 levels following antidepressant treatment ^39 40^. Furthermore, a recent study that isolated astrocyte nuclei using Cx43 as a marker identified NR3C1, encoding the glucocorticoid receptor (GR), as the top upstream regulator of astrocyte-specific DEGs ^41^, linking astrocyte dysfunction to stress signaling pathways. In line with this, differential motif accessibility showed increased accessibility of the NR3C1 motif in protoplasmic astrocyte. These findings suggest altered stress signalling maybe explained by the downregulation of some astrocyte-specific genes.

GLUL was consistently downregulated across modalities, showing decreased gene expression levels in deep layers. GLUL, which encodes glutamine synthetase, is a key astrocyte-specific enzyme involved in the glutamate–glutamine cycle. Reduced GLUL expression may impair glutamate recycling and disrupt excitatory neurotransmission, potentially contributing to altered neuronal function in MDD ^32, 42^. This may be particularly relevant to the dysfunction we have previous described in deep layer excitatory neurons ^2, 4^.

GPR37L1 was also downregulated across modalities and showed decreased expression in deep layers GPR37L1 has been reported to play a role in astrocyte maturation, for instance, loss of GPR37L1 results in reduced morphological complexity, shorter process length, and decreased expression of mature astrocytic genes ^43^. Downregulation of GPR37L1 in depressed individuals has been found across multiple brain regions ^44^. Animal models of depression have also shown downregulation of gene expression and protein levels in the medial PFC ^44^.

Consistent with this, we found that ligand of GPR37L1, PSAP, was downregulated in deep-layer excitatory neurons. Prosaposin (PSAP) has neuroprotective and glioprotective effects such as protection against oxidative stress and cellular damage ^45^. PSAP localizes at the neuron–glia interface to interact with GPR37L1 ^46^. PSAP–GPR37L1 signaling regulates astrocyte homeostasis and neuron–glia interactions. Activation of GPR37L1 by PSAP modulates astrocyte glutamate handling and neuronal NMDA receptor responses ^47^. Furthermore, loss of GPR37L1 increases neuronal vulnerability in models of ischemic injury ^47^, highlighting the importance of PSAP–GPR37L1 signaling in maintaining neuron–glia homeostasis.

When analysing the colocalization of ligands and receptors, we identified Factor 3 as an astrocyte-associated signaling program. This factor was enriched for genes related to astrocyte-related processes, and many of the contributing receptors, including GPR37L1, were predominantly expressed in astrocytes. However, several of the top contributing ligands, including PSAP, GNAS, and CALM1, are primarily expressed in neurons, suggesting that Factor 3 may reflect astrocyte–neuron communication. Indeed, deep-layer excitatory neurons were among the most affected cell populations in MDD in our previous snRNA-seq and snATAC-seq analyses ^2, 4^. Given the critical role of astrocytes in supporting excitatory neurons through metabolic support, glutamate homeostasis, and synaptic regulation, these findings suggest that astrocyte dysfunction may contribute to the communication and vulnerability of excitatory neurons in MDD.

Several studies have examined the role of GPR37L1 in emotional behavior and stress-response regulation in a sex-specific manner. Global knockout of GPR37L1 produces sex-specific changes in anxiety-like behaviors ^48^. Furthermore, astrocyte-specific knockout of GPR37L1 in the hippocampus resulted in trends toward increased anxiety-like behavior in male but not female mice accompanied by reduced plasma corticosterone levels, suggesting sex-specific effect of GPR37L1 loss ^49^.

Consistent with these results, our snRNA data results suggest that astrocyte dysregulation may be more prominent in males compared to females. Indeed, GPR37L1 was differentially expressed in males (adj. p-value = 0.00643) but not in females (adj. p-value = 1). Sex-specific differences were also identified in cell-cell interaction analysis, where PSAP-GPR37L1 signaling was absent in male MDD but reduced in female MDD. However, further analysis is required to elucidate sex-specific dysfunction of these astrocyte specific genes.

There are several limitations to note in our study. First, we present MDD associated molecular changes identified by comparing individuals with MDD to psychiatrically healthy controls, however, all individuals with MDD included in this cohort died by suicide. As a result, it is difficult to distinguish molecular changes associated with MDD from those related from suicide, as these factors are confounded in the present study.

Second, snRNA-seq and snATAC-seq were generated from separate nuclei rather than the same nuclei, although the majority of subjects overlapped between two modalities. Spatial transcriptomics data were also incorporated in a separate batch. A multiome approach, measuring paired gene expression and chromatin accessibility from the same nucleus, would provide greater resolution. Despite these limitations, high concordance in gene expression and gene activities was observed, supporting the robustness of our findings.

In addition, we identified Factor 3, an astrocyte-driven communication program, which showed a trend toward enrichment in MDD. Despite exhibiting one of the strongest enrichment signals observed, the association did not reach statistical significance, likely reflecting limited statistical power. Future studies in larger cohorts will be required to determine whether this trend represents a true biological effect.

Lastly, the resolution of Visium technology limits cell-type-specific interpretation of spatial transcriptomic data. As each spot can contain multiple cells, the measured gene expression represents a mixture of cells rather than a single cell or cell type. Although we identified disease-associated changes in genes known to be predominantly expressed in astrocytes, these findings should be interpreted with caution. Future studies using higher-resolution spatial transcriptomic approaches will help further elucidate our findings.

We provide a multi-omics characterization of astrocyte abnormalities in the dlPFC in MDD by integrating snRNA-seq ^2,3^, snATAC-seq ^4^, and spatial transcriptomic data. We identified spatially localized astrocyte dysfunction in deep cortical layers and astrocyte-driven signaling programs associated with MDD. Our results revealed a potential link between astrocytes and deep-layer excitatory neurons, supported by convergent evidence across modalities. Although further studies are needed to elucidate the functional consequences of these interactions, our findings provide new insight into how spatially coordinated astrocyte dysfunction may contribute to MDD pathology.

## Data availability

snRNA-seq and processed data are available from the NCBI Gene Expression Omnibus database under accession number GSE213982 and GSE144136. snATAC-seq and processed data are available under accession number GSE246443. Visium data will be uploaded and publicly available upon submission. The reference genome version used is available on the 10x Genomics website (refdata-cellranger-arc-GRCh38-2020-A-2.0.0).

## Acknowledge

C.N. is supported by grants from NSERC (RGPIN-2022-03979), NARSAD (BBRF #31204) and CIHR (PJT183904). GT holds a Tier 1 Canada Research Chair in Major Depressive Disorder and Suicide (CRC-2022-00069) and is supported by grants from the Canadian Institutes of Health Research (CIHR; PJT183903, PJT189993, PJT205924), National Institutes of Health (NIH; R01MH131818), and by the Fonds de recherche du Québec through a Research Centre Grant awarded to the Douglas Research Centre - https://doi.org/10.69777/5230. This project has been made possible by the Canada Brain Research Fund (CBRF), an innovative arrangement between the Government of Canada (through Health Canada), and Brain Canada Foundation, the Douglas Hospital Research Centre and the Douglas Foundation, and in part by funding from the Canada First Research Excellence Fund, awarded to McGill University for the Healthy Brains for Healthy Lives initiative, and from the *Fonds de recherche du Québec* - Santé (FRQS). This research was enabled in part by support provided by Calcul Québec (https://www.calculquebec.ca/en/) and the Digital Research Alliance of Canada (alliancecan.ca).

## Author contributions

H.M. conceptualized, performed the analyses, interpreted the results, and wrote the manuscript. H.R. generated the raw data. S.A. supported data investigation. A.C. supported data interpretation. M.A.D provided wet lab support. N.M contributed to sample procurement. C.N. and G.T. supervised study design, data interpretation, and manuscript preparation, and obtained funding.

## Competing interest statement

The authors declare no competing interests.

## Methods

### snRNA-seq and snATAC-seq

We used previously published snRNA-seq data from a cohort of male subjects and female subjects with MDD and matched psychiatrically healthy controls. (Supplementary Table 1). Preprocessing was performed as previously described. Astrocytes were subsequently subset from the integrated dataset based on broad cell-type annotations for downstream analyses. We also used previously published snATAC-seq data from a cohort with 94% subject overlap with the snRNA-seq dataset (Supplementary Table 1). Astrocyte subtype annotations from the original study were used for all analyses. Differentially accessible region and transcription factor motif enrichment results were retrieved from the original publication.

### snRNA-seq Processing

snRNA-seq data were processed using Seurat ^22^. Subjects with fewer than 20 astrocyte nuclei were excluded from following analyses. SCTransform normalization was performed independently for each subject, with the percentage of mitochondrial transcripts regressed out. Variable features were identified within each sample and integrated across samples using SelectIntegrationFeatures. The sample-level SCTransform-normalized objects were then merged and used for downstream dimensionality reduction. Principal component analysis (PCA) was performed using 50 components. To account for technical variation, Harmony was applied using batch, sequencing chemistry, and subject identity as covariates. The Harmony-corrected embeddings were used to construct a shared nearest-neighbor graph for clustering. To evaluate clustering stability, silhouette scores were calculated across clustering resolutions ranging from 0.1 to 2.0 (Supplementary Figure 2A). A clustering resolution of 0.3 yielded the highest silhouette score and identified five astrocyte clusters, which were used for all downstream analyses. Cluster marker genes were identified using FindAllMarkers with default parameters, comparing each cluster against all remaining cells (Supplementary Table 3).

### MetaNeighbor

MetaNeighbor ^18^ was used to assess the correspondence between astrocyte subtypes identified in the snRNA and snATAC dataset. The snRNA dataset was used as the reference to train the model using defaults parameters. Cell-type similarity was measured using AUROC scores, and the resulting subtype correspondences were visualized using best-hit plots.

### Spatial label transfer

Astrocyte subtypes identified in the snRNA were label transferred to the spatial transcriptomics dataset using FindTransferAnchors in Seurat. Both the snRNA and spatial transcriptomics datasets were normalized using SCTransform. Transfer anchors were identified using the top 50 principal components, and prediction score were computed and plotted.

### Differential gene expression analysis

Pseudobulk differential gene expression analysis was performed using muscat ^50^ and edgeR ^51^ as previously described. Analyses were performed separately in males and females, and results were combined using meta-analysis. Only subjects with a minimum of 5 cells per subtype was kept. Consistent with the original study, Age, brain pH, postmortem interval (PMI), and batch were included as covariates. DEGs were defined using an adj. p-value < 0.1 and |logFC| > log2(1.1). Gene Ontology (GO) enrichment analysis for biological process terms was performed using clusterProfiler ^52^, with p-value < 0.05 were applied as a cut-off.

### snRNA-seq cell-cell interaction analysis

Cell-cell communication analysis between protoplasmic astrocytes and ExN10_L46 excitatory neurons was performed using CellChat following the vignette ^25^. Analyses were conducted separately for males and females, and CellChat objects were constructed for each group using default parameters. Intercellular communication networks were inferred using the computeCommunProb with the trimean method to estimate average gene expression within each cell population. CellChat objects from control and MDD samples were subsequently merged for comparative analysis. Differential signaling interactions were assessed by comparing communication probabilities between conditions. PSAP–GPR37L1 ligand-receptor interaction was extracted and plotted.

### Postmortem brain samples for spatial transcriptomics

Frozen human brain samples dissected from Brodmann area 9, dorsolateral prefrontal cortex (dlPFC), was obtained from the Douglas Brain Bank (RRID:SCR_025991) in collaboration with the Quebec Coroner’s Office. This project was approved by the Douglas Hospital Research Center institutional Ethic review board. Comprehensive psychological autopsies were conducted. Briefly, these consist of a series of proxy-based, structured interviews assessing psychopathology with next-of-kin and reviews of medical records. Cases were individuals who died by suicide in the context of an episode of MDD. Controls were individuals who died suddenly and did not have evidence of an axis I disorder. Group was matched for postmortem interval (PMI), tissue pH, and RNA integrity number (Supplementary Table 1).

### Spatial transcriptomics library preparation and sequencing

Spatial transcriptomic profiles were generated using the 10x Genomics Visium Spatial Gene Expression platform (version 2). Frozen, unfixed brain tissue blocks were mounted onto specimen chucks using M-1 Embedding Matrix (Epredia) and cryosectioned at −21°C. Prior to sectioning, the face of each tissue block was trimmed with a single-edge razor blade to approximately 6.5 mm^2^ to fit within the fiducial frame of each Visium capture area. A single 10 µm tissue section from each specimen was collected and placed onto a capture area of the slide. Slides were immediately stored at −80°C until further processing. Subsequent tissue processing and library preparation were performed according to the Visium Spatial Gene Expression User Guide (Rev F, CG000239). Final dual-indexed libraries were sequenced on an Illumina NovaSeq 6000 system using a NovaSeq S4 reagent kit. Sequencing was performed using the following read configuration: Read 1, 28 cycles; i7 index, 10 cycles; i5 index, 10 cycles; and Read 2, 91 cycles.

### Spatial transcriptomics preprocessing

Sequencing data were processed using Spaceranger (10X Genomics) with default parameters. Reads were aligned to the GRCh38 reference genome (GRCh38-2020-A) and gene expression matrices were generated. Quality metrics generated by Spaceranger is provided in Supplementary Table 2. Spots were further removed based on the following criteria. Spot with zero count and outside the tissue were removed. Spots with high mitochondrial expression (>2 MADs above the sample-specific median), low library size (<3 MADs below the median), or low numbers of detected genes (<3 MADs below the median) were removed, resulting in the exclusion of 4,472 spots (8.0%) (Supplementary Figure 4-6). One sample was subsequently excluded due to high mitochondrial content. The final dataset consisted of 48,673 high-quality spots for downstream analyses.

### Spatial domain clustering

Gene expression of marker of neurons, astrocytes, and deep layer showed cortical layer organization (Supplementary Figure 7-9). To identify cortical layer spatial domains, unsupervised clustering was performed using BANKSY ^19^. The top 2,000 highly variable genes were identified for each sample, and gene counts were normalized. BANKSY neighborhood matrices were computed with kgeom =18, corresponding to first- and second-order neighboring spots. PCA was performed using 10 PCs with a spatial weighting parameter λ=0.2. Harmony was performed to account for batch effects across samples with max.iter.harmony = 20. Clustering resolution parameters ranging from 0.1 to 2.0 were evaluated, and a resolution of 0.6 was selected as the primary resolution based on correspondence with known cortical laminar structures and cluster stability (Supplementary Figure 10B). Clustering stability was assessed using the FAST+ algorithm (Supplementary Figure 10A). Spatial domains representing fewer than 0.1% of all spots were excluded from downstream analyses.

### Spatial domain annotation

To annotate spatial domains to cortical layers and white matter, a publicly available human dlPFC dataset with manual layer annotations was used as a reference following a previously published approach using spatialLIBD ^20^. Briefly, gene expression profiles were aggregated across spots within each spatial domain to generate pseudobulk expression matrices. Differentially expressed genes were identified for each domain, and domain-specific enrichment scores were correlated with those derived from the reference dataset to assign cortical layer identities (Supplementar Table 5). Based on this approach, spatial domains were annotated as c1-WM, c2-WM/L6, c3-L6, c4-L5, c5-L4, c6-L3, and c7-L2.

### Cell type deconvolution using cell2location

Cell type abundance in each spot was estimated using cell2location ^21^. The snRNA-seq dataset was used as a reference to infer cell-type signatures. Cell-type deconvolution was performed at two levels: (i) broad cell-type annotations and (ii) fine-resolution annotations incorporating excitatory neuron subtypes with layer-specific identities. Cells annotated as “Mix” were excluded from the reference dataset. Reference cell-type signatures were estimated using the negative binomial regression model implemented in cell2location. The trained reference signatures were subsequently used to estimate the abundance of each cell type within individual Visium spots by deconvolving spot-level gene expression profiles. Following a previous study of human dlPFC Visium data, the expected cell abundance was set to five cells per spot, based on manual histological quantification.

### Layer-specific pseudobulk differential expression analysis

Pseudobulk profiles were generated by aggregating counts across spots from the same subject within each spatial domain using registration_pseudobulk, with a minimum of 10 spots required per subject-domain pair. Differential expression analysis was performed separately for each spatial domain using the limma-voom workflow ^53^. Lowly expressed genes were filtered using filterByExpr, and normalization factors were calculated using edgeR. Disease status was modeled as the primary variable of interest, with slide ID and brain pH included as covariates. DEGs were defined using two significance thresholds: (i) adj. p-value < 0.1 and |FC| > 1.5, and (ii) nominal p-value < 0.05 and |FC| > 1.5. GO enrichment analysis for biological process terms was performed using clusterProfiler ^52^, with p-value < 0.05 were applied as a cut-off.

### Ligand–receptor spatial co-expression analysis

LIANA+ was used to identify spatially co-expressed ligand–receptor pairs across samples ^24^. Spatial neighbors were defined using spatial neighbors with a bandwidth= 200, corresponding to first-order neighboring spots. For each ligand–receptor pair, a spatial co-expression score was computed using weighted cosine similarity. Ligand–receptor pairs expressed in at least 10% of spots were retained for downstream analysis. Non-negative matrix factorization (NMF) was then applied to identify spatial ligand–receptor co-expression programs across slides. The optimal number of programs was determined using the elbow method with default parameters, resulting in five programs. Mann-Whitney was used to assess enrichment of specific programs between groups, based on spots with average factor scores above the 75th percentile. For each program, ligand–receptor pairs with the highest factor loadings were considered the top contributing interactions. Functional enrichment analysis of the top contributing ligand and receptor genes was performed using the clusterProfiler ^52^, using default parameter with p-value < 0.05 as a cut-off.

## Notes

### Competing Interest Statement

The authors have declared no competing interest.

