## Supplemental Figure for "Multi-omics characterization of astrocyte subtypes reveals spatially coordinated astrocyte downregulation in depression"

#### Supplementary Figure 1

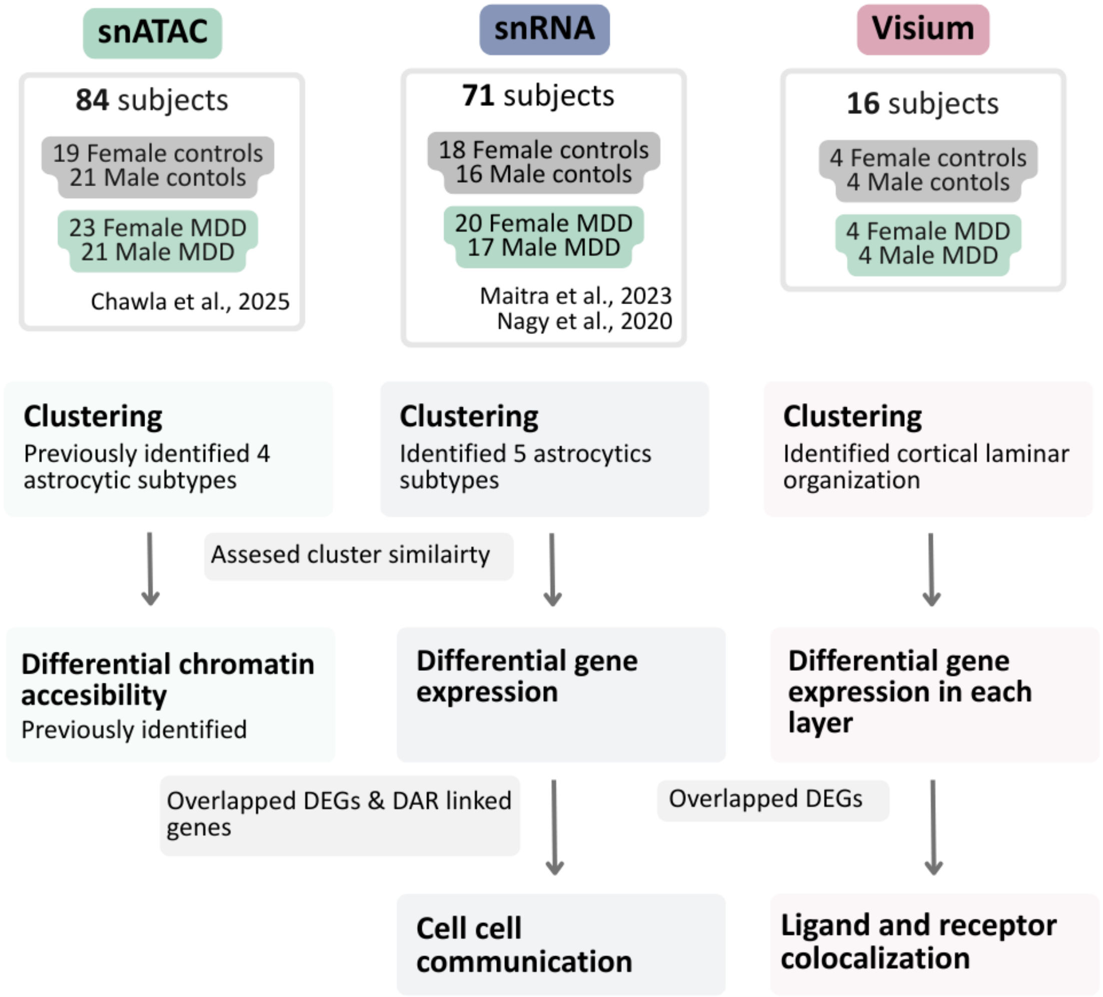


**Supplementary Figure1** Schematic overview of the study design. snATAC-seq, snRNA-seq, and Visium spatial transcriptomic datasets from control and MDD subjects were integrated to characterize astrocyte subtypes, cortical layer organization, differential chromatin accessibility, differential gene expression, cell–cell communication, and ligand–receptor colocalization.

#### Supplementary Figure 2

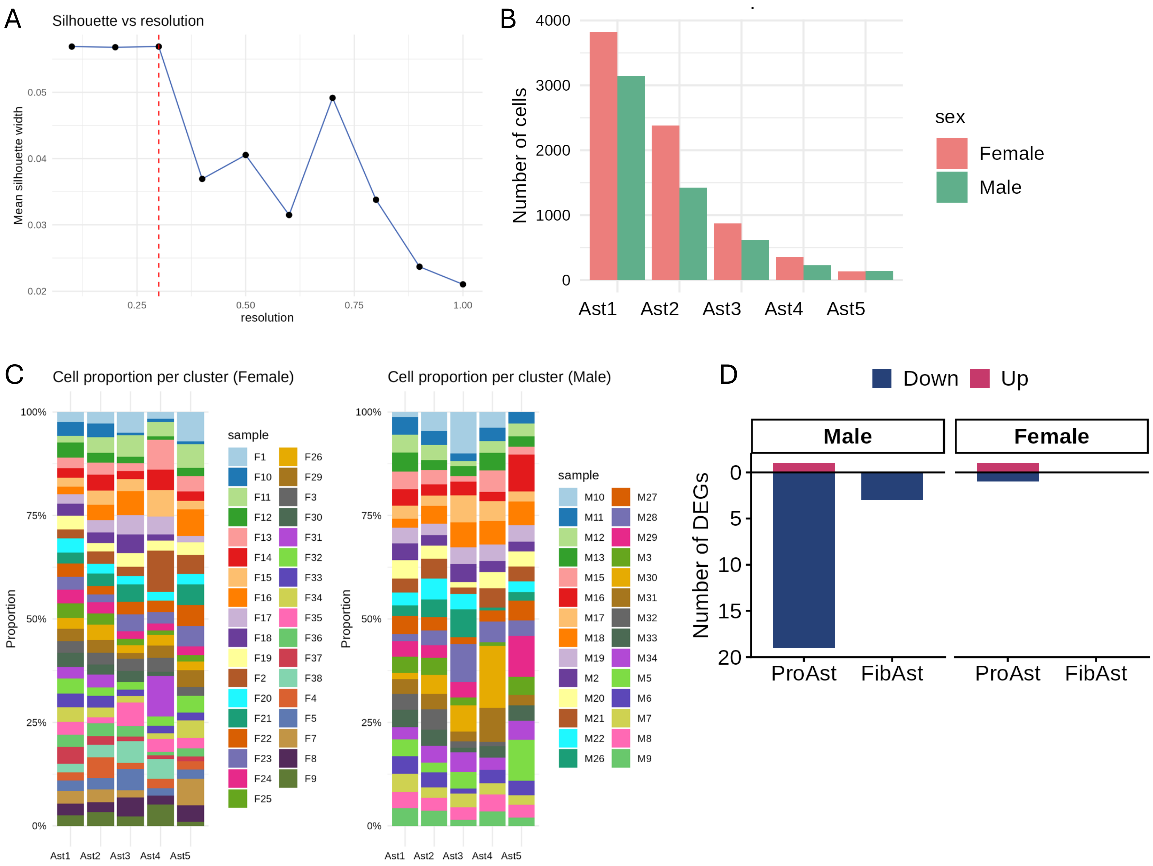


**Supplementary Figure2 A.** Clustering resolution for snRNA-seq astrocyte subclusters was determined based on clustering stability measured by silhouette score. Clustering stabilized at resolution 0.3 (red line), which was selected as the final clustering resolution. **B.** Number of nuclei in each astrocyte subcluster, shown separately for females and males, as the datasets were generated in separate batches. **C.** Proportion of nuclei contributed by each sample to each astrocyte subcluster, shown for females (left) and males (right). **D.** Number of DEGs identified in each astrocyte subtype in MDD compared to controls in each sex. Bar is colored by direction of fold change (downregulated = blue, upregulated = red), using a threshold of adjusted p-value < 0.1 and |logFC| > log2(1.1).

### Supplementary Figure 3

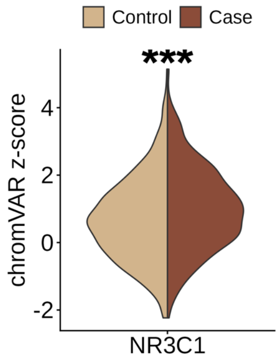


**Supplementary Figure3** Violin plots showing chromVAR deviation (z-score) for the NR3C1 motif in Ast1 from control and MDD subjects (adj. p = $7.28 \times 10^{-5}$).

#### Supplementary Figure 4

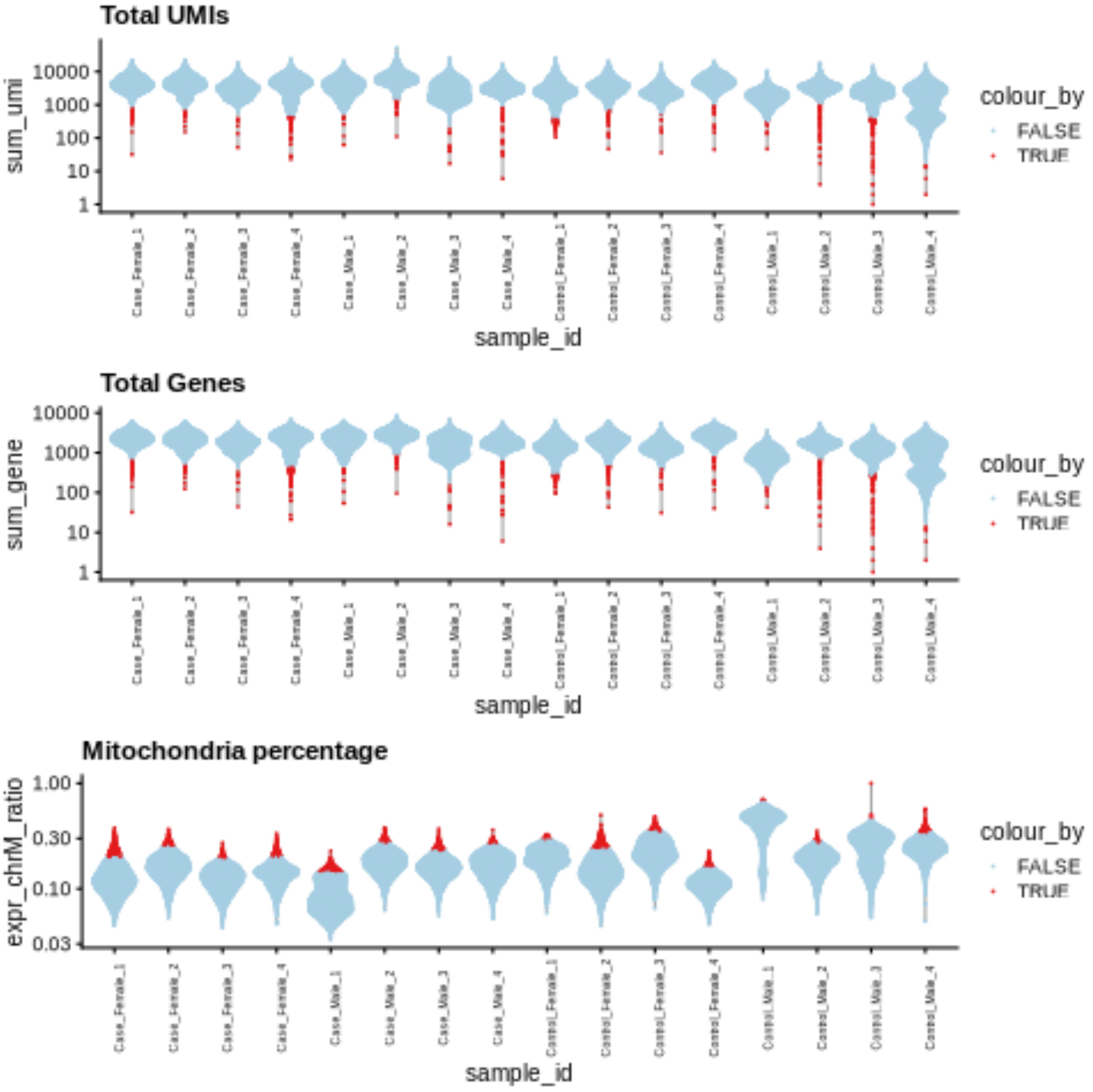


**Supplementary Figure4** Spots were filtered based on three quality metrics: total UMI count, total number of detected features, and mitochondrial gene percentage. Outlier spots were identified using median absolute deviation (MAD)-based thresholds. Spots retained for downstream analysis are shown in blue and outlier spots removed from analysis are shown in red.

### Supplementary Figure 5

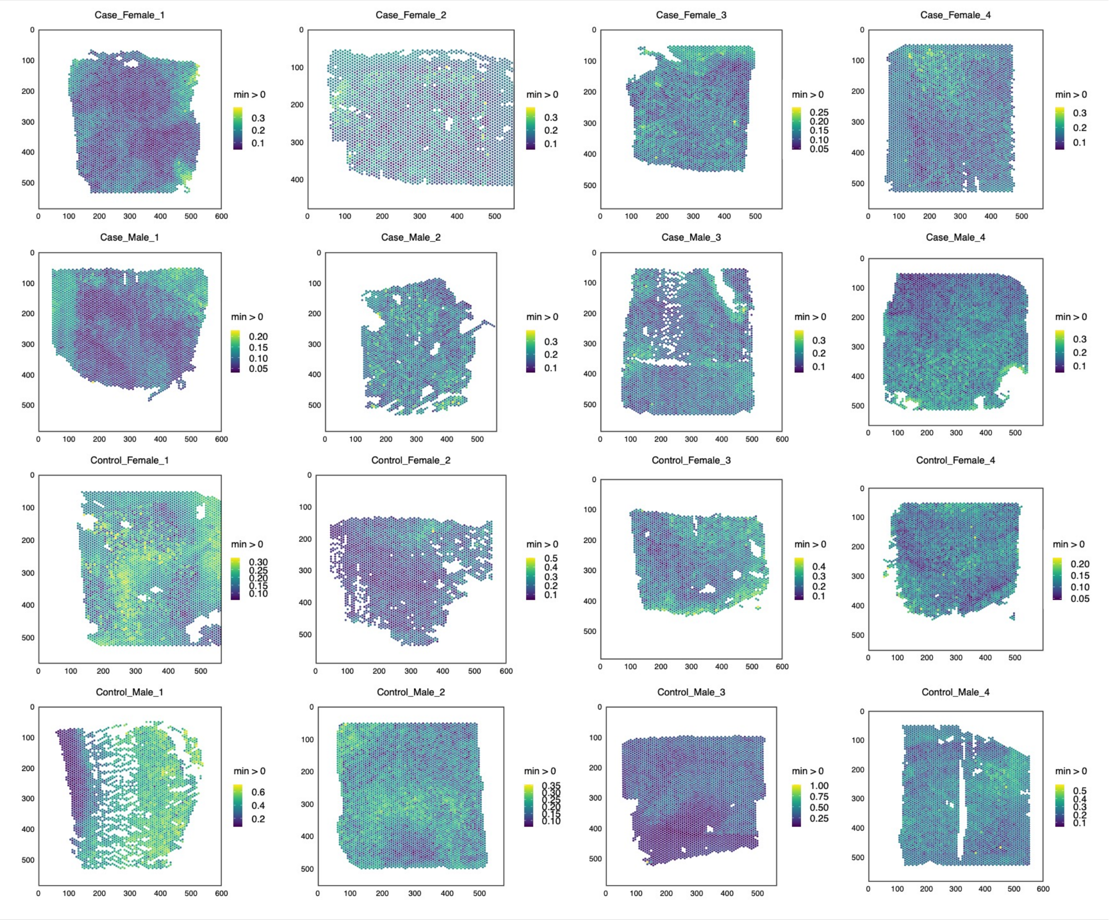


**Supplementary Figure5** Feature plot showing mitochondrial gene percentage across Visium spots, reflecting the laminar organization of the dlPFC.

### Supplementary Figure 6

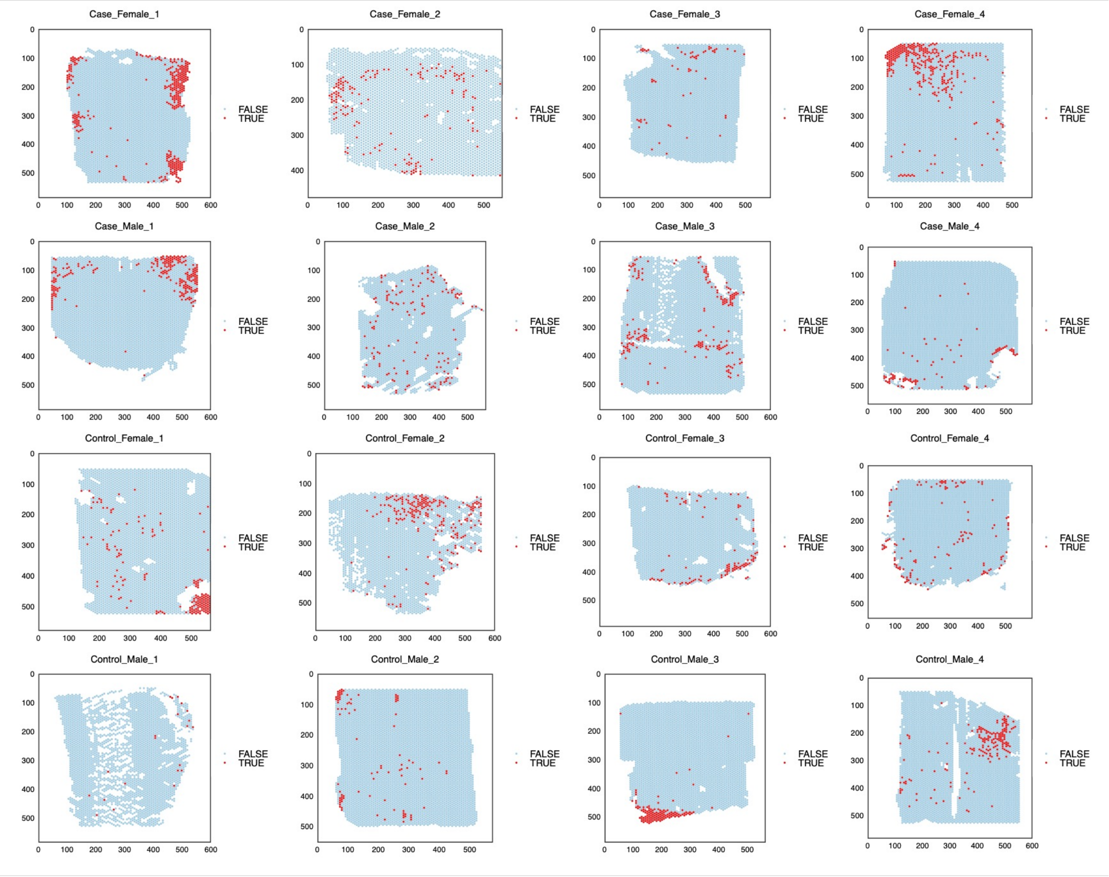


**Supplementary Figure6** Feature plots showing discarded spot across Visium spots. Spots retained for downstream analysis are shown in blue and outlier spots removed from analysis are shown in red.

### Supplementary Figure 7

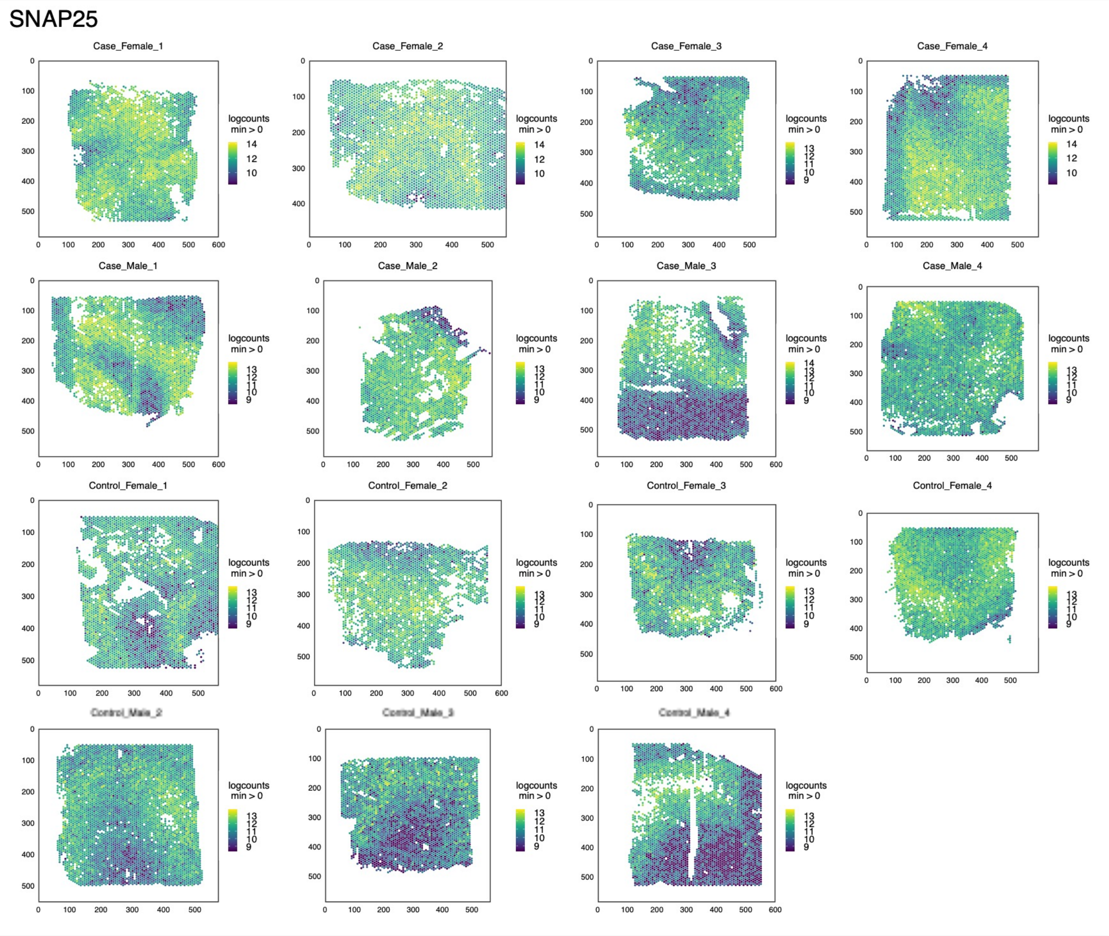


**Supplementary Figure7** Feature plots showing SNAP25, a neuronal marker, across Visium spots in each subject. Showing distribution in cortical layers.

#### Supplementary Figure 8

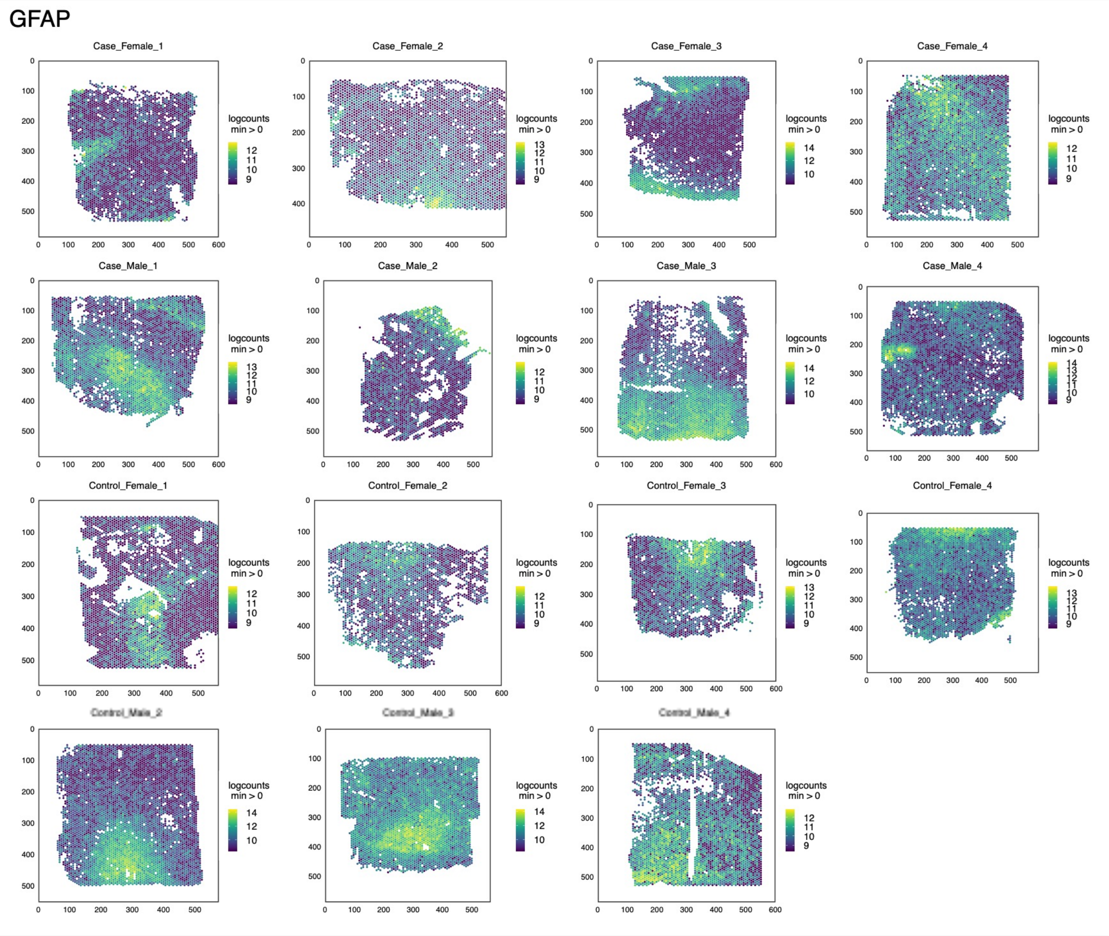


**Supplementary Figure8** Feature plot showing GFAP, an astrocytic marker, across Visium spots in each subject. Showing distribution in white matter.

### Supplementary Figure 9

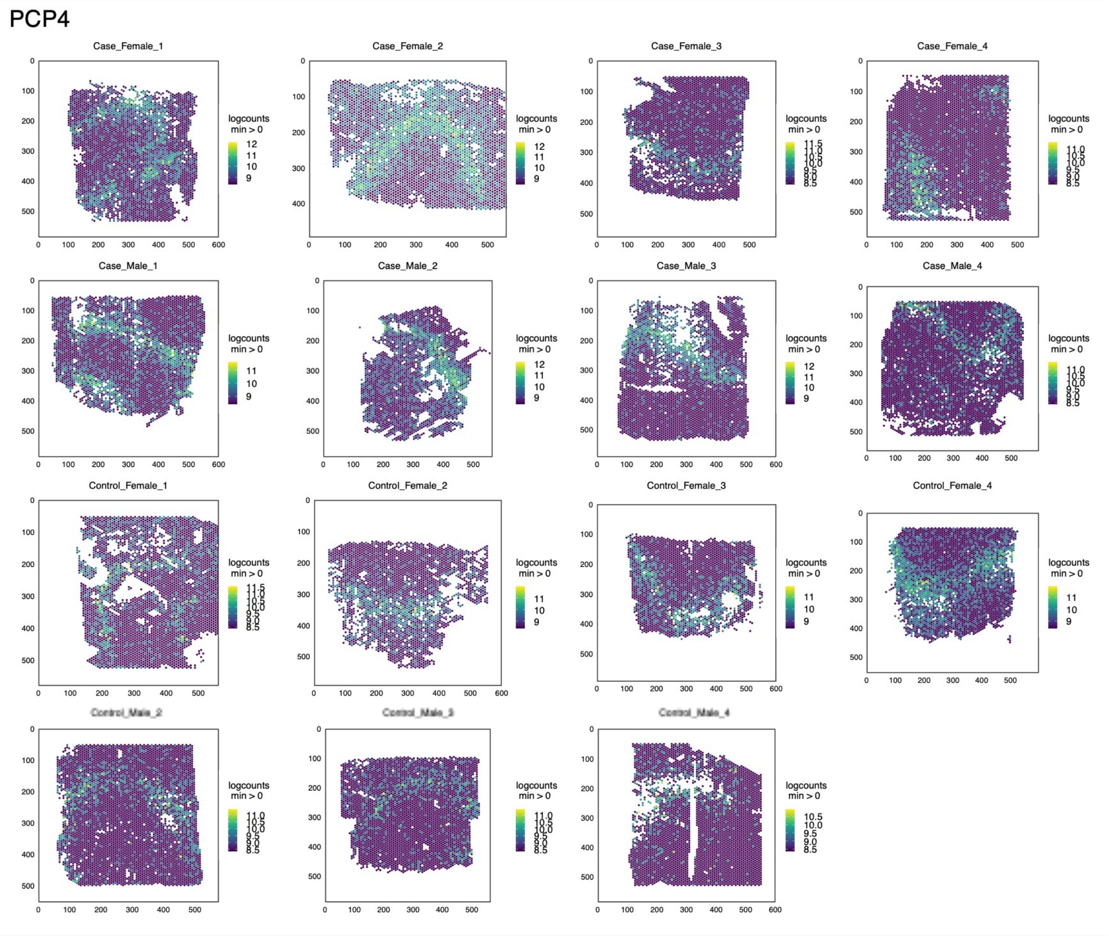


**Supplementary Figure9** Feature plots showing PCP4, a deep layer excitatory marker, across Visium spots in each subject. Showing distribution in deep cortical layers.

### Supplementary Figure 10

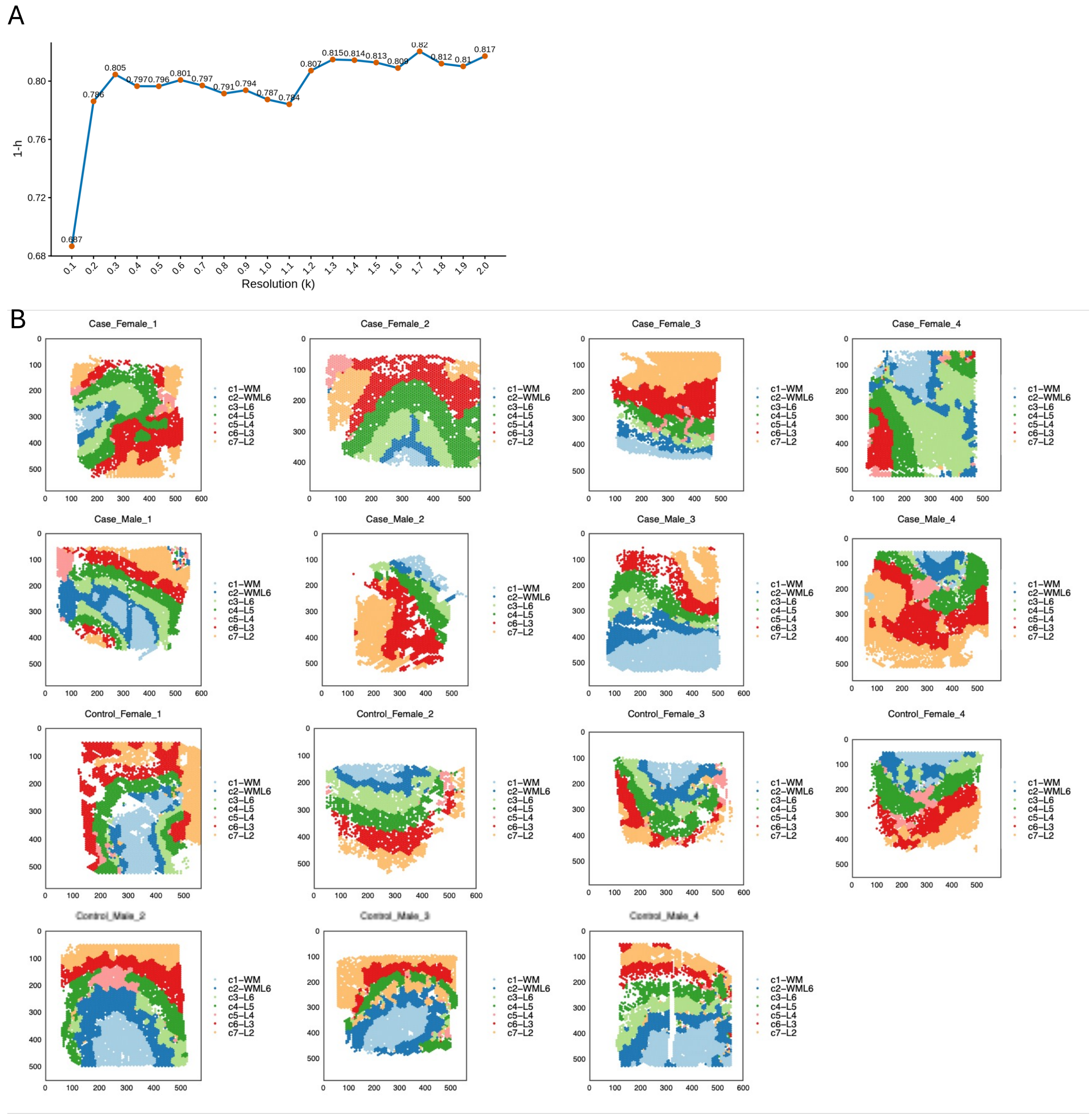


**Supplementary Figure10 A.** Determination of clustering resolution for cortical layer identification using FastHPlus. Clustering stability was evaluated across resolutions ranging from 0.1 to 2.0 using the 1 − h score. The final resolution of 0.6 was selected based on both clustering stability and agreement with the known laminar organization of the cortex, consisting of six cortical layers and white matter. **B.** Feature plot showing clustering results colored by annotated cortical layers in each subject.

### Supplementary Figure 11

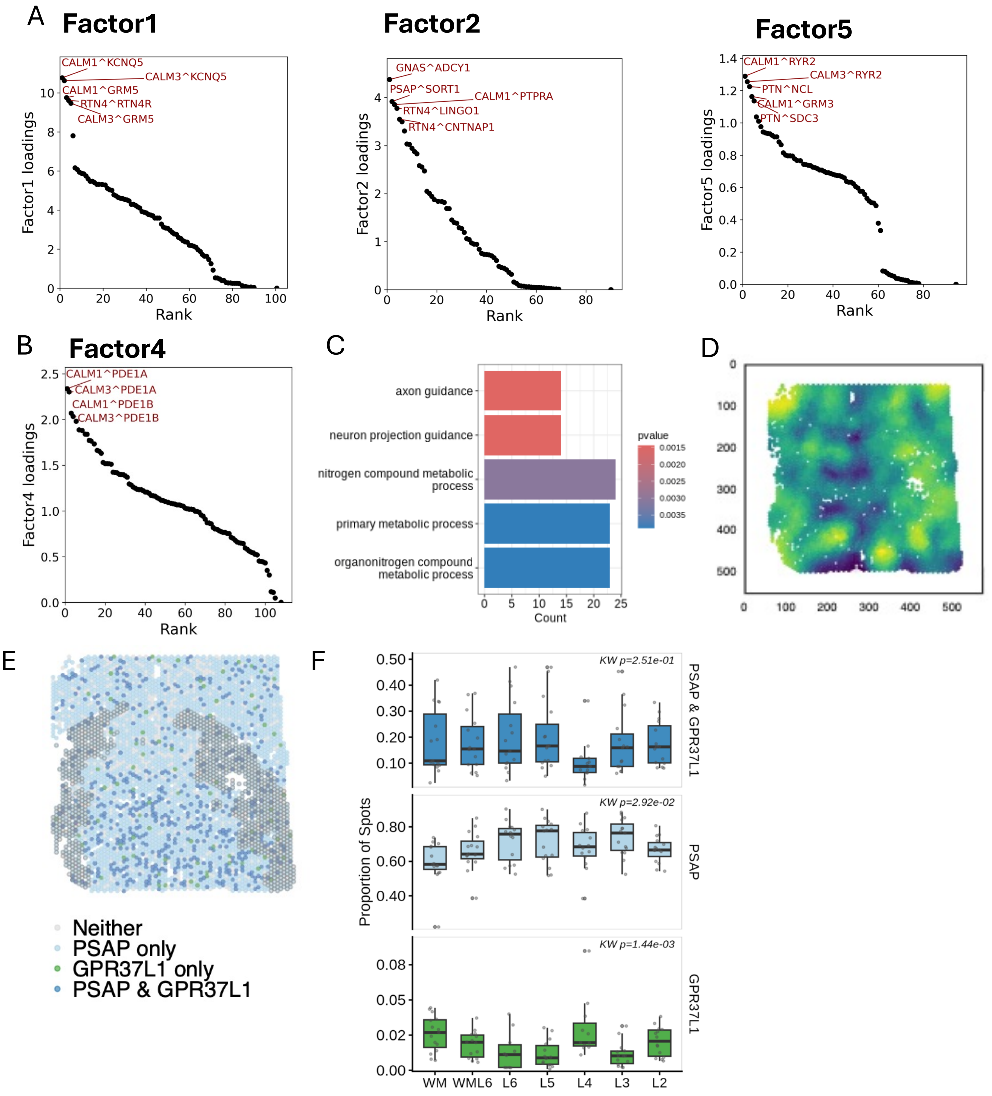


**Supplementary Figure11 A.** Ranked contribution of ligand-receptor pairs to Factor 1, 2, and 5 loading scores, showing the top 5 contributing pairs. **B.** Ranked contribution of ligand-receptor pairs to Factor 4 loading scores, showing the top 5 contributing pairs. **C.** Gene ontology enrichment analysis of the top contributing ligands and receptors in Factor 4. **D.** Spatial feature plots showing local co-expression scores of CALM1 and PDE1A across spots. **E.** Spatial distribution of PSAP and GPR37L1 expression in a representative sample. Light blue spots indicate PSAP-only expression, green spots indicate GPR37L1-only expression, and dark blue spots indicate co-expression of PSAP and GPR37L1. Spots outlined in black correspond to deep cortical layers (L5 and L6), which were highlighted because GPR37L1 was downregulated in cases in these layers. **F.** Distribution of PSAP- and GPR37L1-expressing spots across cortical layers. Bar plots show the average proportion of co-expressing (PSAP+/GPR37L1+), PSAP-only, and GPR37L1-only spots across all samples and cortical layers. Differences in spatial distributions across layers were assessed using the Kruskal–Wallis test.
